# Defective synapse maturation and enhanced synaptic plasticity in Shank2^-/-^ mice

**DOI:** 10.1101/193078

**Authors:** Stephanie Wegener, Arne Buschler, A. Vanessa Stempel, Sarah A. Shoichet, Denise Manahan-Vaughan, Dietmar Schmitz

**Affiliations:** Neuroscience Research Center, Charité Universitätsmedizin, Berlin, Germany; Present address: Janelia Research Campus, Howard Hughes Medical Institute, Ashburn VA 20147, USA; Ruhr University Bochum, Universitätsstr. 150, Bochum, Germany; Present address: Sainsbury Wellcome Centre, 25 Howland Street, London, United Kingdom; Institute of Biochemistry and Cluster of Excellence NeuroCure, Charité Universitätsmedizin, Berlin, Germany; Neuroscience Research Center,Cluster of Excellence NeuroCure, German Center for Neurodegenerative Diseases,Charité Universitätsmedizin, Berlin, Germany

**Author notes:** Corresponding authors: Stephanie Wegener Dietmar Schmitz.

## Abstract

Autism spectrum disorders (ASDs) are neurodevelopmental disorders with a strong genetic aetiology. Since mutations in human SHANK genes have been found in patients with autism, genetic mouse models are employed for a mechanistic understanding of ASDs and the development of therapeutic strategies. In sharp contrast to all studies so far on the function of SHANK proteins, we observe enhanced synaptic plasticity in Shank2^-/-^ mice, under various conditions *in vitro* and *in vivo*. Reproducing and extending previous results, we here present a plausible mechanistic explanation for the mutants' increased capacity for long-term potentiation (LTP) by describing a synaptic maturation deficit in Shank2^-/-^ mice.

## Introduction

The activity-dependent formation and remodelling of synaptic connections is pivotal to adaptive neural circuit function. Dysregulation of these processes is considered a prime cause of neurodevelopmental diseases such as autism spectrum disorders (ASDs) (Ebert and Greenberg, 2013). The list of mutations associated with ASDs and the number of mouse models is growing rapidly, yet understanding the connection between genetic mutation, synaptic defects, and disease phenotypes remains a challenge. Mutations in *SHANK* genes occur in autistic patients (Durand *et al.*, 2007; Berkel *et al.*, 2010; Sato *et al.*, 2012) and Shank mutant mice reproduce several autism-related phenotypes (Bozdagi *et al.*, 2010; Peça *et al.*, 2011; Wang *et al.*, 2011; Schmeisser *et al.*, 2012; Won *et al.*, 2012; Lee *et al.*, 2015; Peter *et al.*, 2016). SHANKs (short for SH3 and multiple ankyrin repeat domains protein, also referred to as ProSAPs) are scaffold proteins in the postsynaptic density of mammalian excitatory synapses that interact with numerous synaptic proteins (Monteiro and Feng, 2017). Recent reports have made a strong case for NMDA receptor hypofunction and/or defective long-term potentiation as a cause for disease-relevant behavioural phenotypes in Shank2 and Shank3 mutant mice (Won *et al.*, 2012; Lee *et al.*, 2015; Peter *et al.*, 2016) and Shank3 (Bozdagi *et al.*, 2010; Peça *et al.*, 2011; Kouser *et al.*, 2013) and it has even been suggested that pharmacological upregulation of LTP can be used to reverse the phenotype (Won *et al.*, 2012; Bozdagi, Tavassoli and Buxbaum, 2013). In sharp contrast to all other studies on the function of SHANK proteins we observe an *enhancement* of synaptic plasticity in Shank2^-/-^ mice (Schmeisser *et al.*, 2012). Here, we report robustly increased LTP *in vitro* and also following *in vivo* induction of LTP in Shank2^-/-^ mice. Moreover, we find that Shank2^-/-^ mice suffer from deficient synaptic maturation, suggesting a mechanistic explanation for their increased LTP capacity. This calls for further investigations into how autism-associated phenotypes reflect alterations in synaptic maturation and stability before we should attempt to pharmacologically increase LTP in autism therapies.

## Results and Discussion

Motivated by methodological differences between our previous findings (Schmeisser *et al.*, 2012) and a parallel study (Won *et al.*, 2012), that reported increased (Schmeisser *et al.*, 2012) versus decreased LTP (Won *et al.*, 2012), respectively, in genetically similar Shank2^-/-^ mice, we first investigated whether differences in slice storage, recording temperature, and animal age might account for the discrepancy in the expression of LTP. To this end, we reproduced a number of conditions from Won et al. (2012), i.e. slices were stored in an ACSF/oxygenated air (Haas-type) interface chamber instead of submerged in ACSF, animal age was 8-9 instead of 3-4 weeks, and recordings were performed at elevated instead of room temperature. Hippocampal slices from Shank2^-/-^ mice were then subjected to the common LTP induction protocol (see Methods for details). In our hands, also in these conditions, Shank2^-/-^ mice showed markedly higher LTP than wild-type controls (Figure 1A). Slice storage seemed to affect different electrophysiological phenotypes to various degrees, however. While storing slices submerged in ACSF versus a Haas-type interface chamber had no effect on the expression of LTP (Figure 1B) or basal synaptic transmission (Figure 1C), it did affect the observed AMPA/NMDA receptor ratios (Figure 1D).

**Figure 1.**
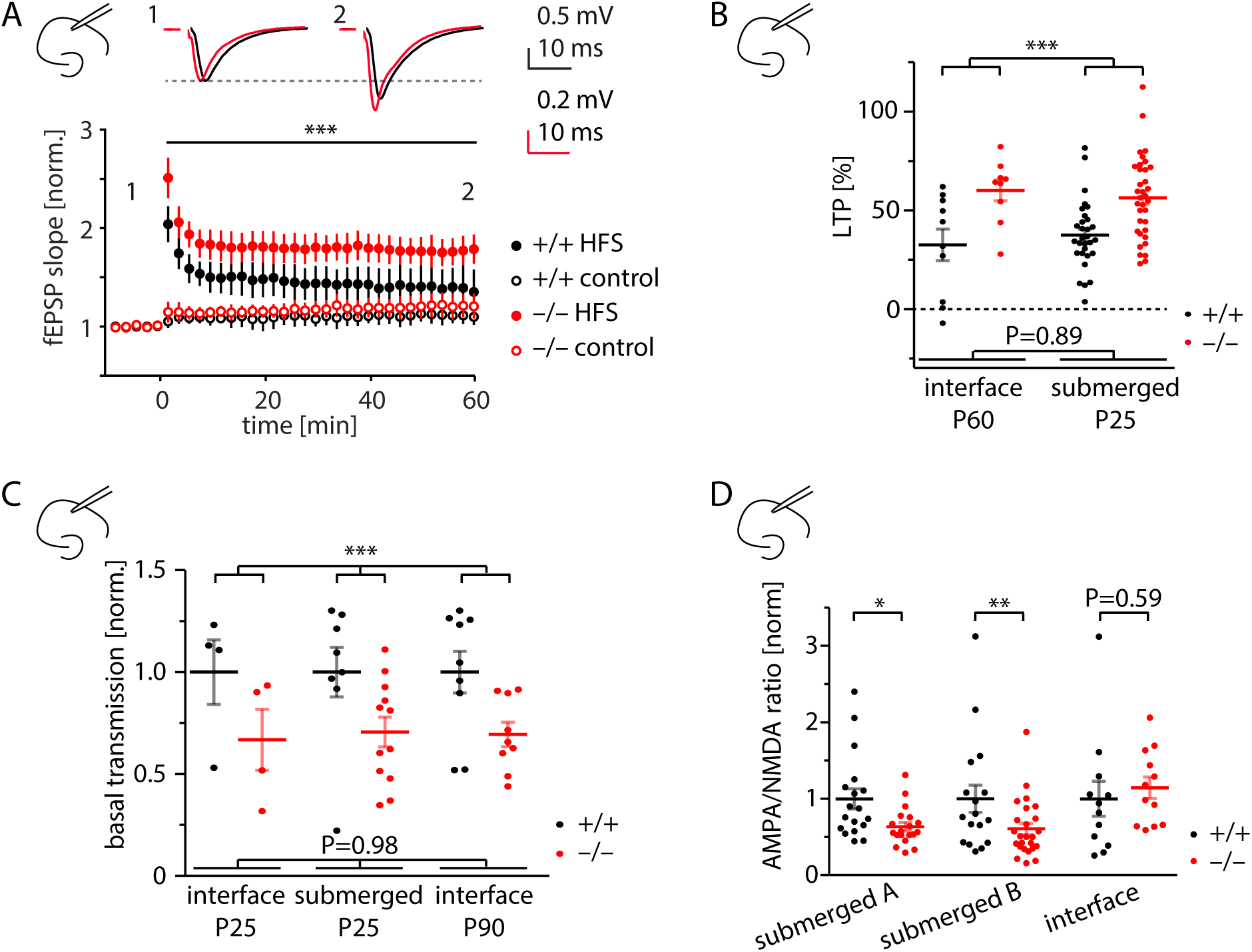
Enhanced LTP and reduced basal synaptic transmission in Shank2^−**/**–^ **mice *in vitro***. A. Enhanced long-term potentiation in Shank2^−/–^ mice *in vitro*. NMDA receptor dependent LTP in the CA1 region of corticohippocampal slices was induced by high frequency stimulation protocol (‘HFS’, closed symbols). In a subset of experiments (+/+: 5/10, –/–: 7/9), synaptic responses of a non-potentiated fiber tract were recorded as an additional control (‘control’, open symbols). Example traces at top. **: p=0.0025 (difference between genotypes in a two-way ANOVA; n(N)_+/+_ = 10(3), n(N)_–/–_ = 9(3)). Slices were stored in an ACSF/oxygenized air interface chamber as reported by Won et al. (2012). B. Increased LTP in Shank2^−/–^ mice is irrespective of animal age and slice storage. Submerged data replotted from Schmeisser et al. (2012). Significance was tested with two-way ANOVA (***: p<0.0001 for genotype comparison across conditions; n(N)_+/+ interface_ = 10(3), n(N)_–/–interface_ = 9(3), n(N)_+/+ submerged_ = 30(5), n(N)_–/–submerged_ = 34(6)). C. Decreased basal synaptic transmission in Shank2^−/–^ mice irrespective of animal age and slice storage. Basal transmission in each experiment is expressed as a single slope fitted to the input-output function of fEPSP slope versus fibre volley. Slopes of each group are normalized to the population mean slope of the wildtype in the respective recording condition. Submerged data (P25) and interface data (P90) are re-analyzed from Schmeisser et al. (2012). Significance was tested with two-way ANOVA (***: p=0.0005 for genotype comparison across conditions; n(N)_+/+ interface P25_ = 4(2), n(N)_–/–interface P25_ = 4(2), n(N)_+/+ submerged P25_ = 8(3), n(N)_–_/– submerged P25 = 11(4), n(N)_+/+ interface P90_ = 9(3), n(N)_–/–interface P90_ = 9(3)). D. Slice storage conditions affect AMPA/NMDA receptor ratios. Significance was tested with Student’s t-test. AMPA/NMDA receptor ratios are significantly reduced in CA1 pyramidal cells of Shank2^−/–^ mice when slices are stored submerged in ACSF (‘submerged 1’: p = 0.013 (*), data replotted from Schmeisser et al. (2012), n(N)_+/+_ = 18(8), n(N)_–/–_ = 19(6); ‘submerged 2’: p = 0.0012 (**), n(N)_+/+_ = 20(6), n(N)_–/–_ = 18(8), see also Figure 3C), but not when stored in an ACSF/oxygenated air interface chamber (‘interface’: n(N)_+/+_ = 12(2), n(N)_–/–_ = 12(2)). Mouse age was 3-4 weeks for all groups.

In light of these results and reports on substantial changes in synaptic spine morphology following brain slice preparation (Kirov, Sorra and Harris, 1999), we reasoned that an *in vitro* examination of Shank2^-/-^ phenotypes might be problematic, given that SHANK2 has a well-described role in synaptogenesis and the regulation of structural dynamics in dendritic spines (MacGillavry *et al.*, 2016). In particular we wondered whether the decreased basal synaptic transmission and increased LTP we consistently observed in Shank2^-/-^ mice might be secondary to slicing-induced synaptic remodelling. This motivated us to examine synaptic transmission and LTP *in vivo*. Most rodent *in vivo* LTP studies have been performed in rats rather than mice, however, and not all protocols that generate LTP in brain slices consistently generate lasting synaptic plasticity *in vivo*, possibly due to differences in neural dynamics and neuromodulatory tone in the intact brain. Using recently established experimental procedures (Kemp and Manahan-Vaughan, 2008) (see Methods for details), we compared Shank2^-/-^ and wild-type mice with regard to their capacity to express electrically induced LTP *in vivo*. We tested different induction protocols, eliciting both short-and long-lasting forms of LTP (Kemp and Manahan-Vaughan, 2008). Both forms of LTP could be elicited in Shank2^-/-^ mice and wildtype controls (Figure 2A, B), validating that synapses without SHANK2 can express LTP, as our data from acute slices suggest. For long-lasting LTP, the potentiation in Shank2^-/-^ mice significantly exceeded that of wild-type controls (Figure 2A). Further corroborating our *in vitro* results, basal synaptic transmission was significantly decreased in Shank2^-/-^ mice versus wildtype controls when assessed in awake, behaving animals (Figure 2C). In summary, we observe enhanced LTP and decreased basal synaptic transmission in Shank2^-/-^ mice under a range of different conditions both *in vitro* and *in vivo*.

**Figure 2.**
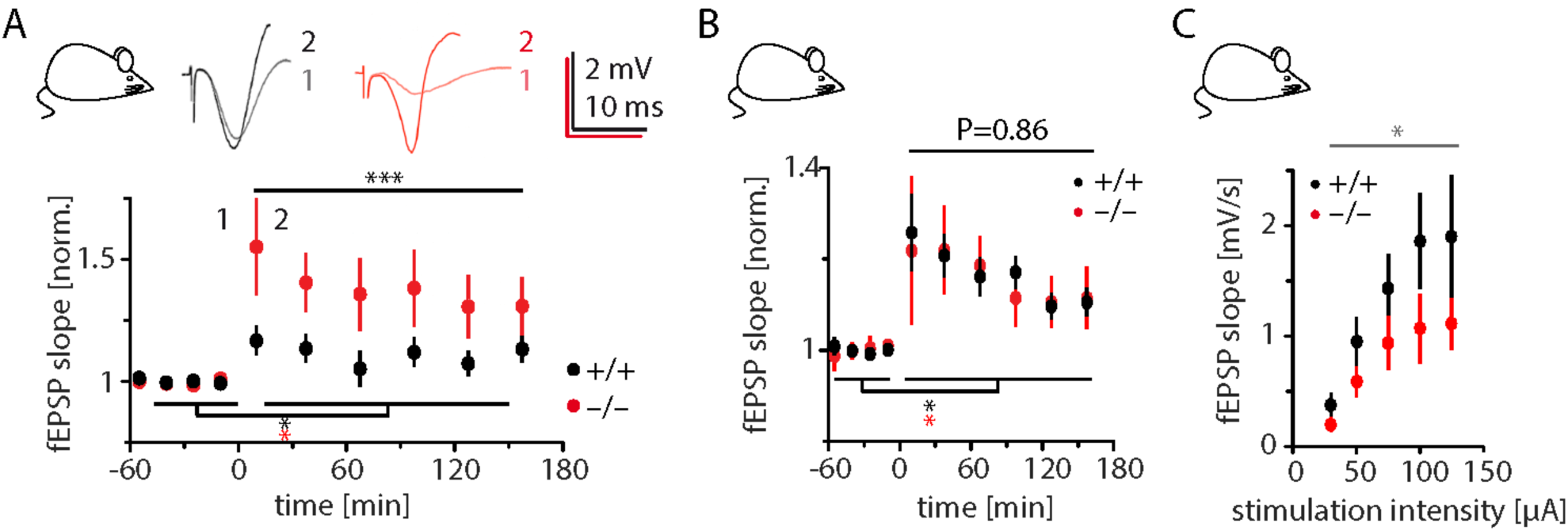
Enhanced LTP and reduced basal synaptic transmission in Shank2^−/–^ mice *in vivo*. A. Enhanced long-term potentiation in Shank2^−/–^ mice *in vivo*. NMDA receptor dependent LTP was induced by strong high frequency stimulation in awake, behaving mice (see Methods for details). Example traces at top. LTP was induced in both genotypes as tested by ANOVA (*: +/+: p<0.0001, N = 16; –/–: p=0.0125, N = 15.) but was significantly larger in Shank2^−/–^ mice (***: p<0.0001, difference between genotypes in a two-way ANOVA). B. NMDA receptor dependent LTP was induced by mild high frequency stimulation in awake, behaving mice (see Methods for details). Significance was tested with two-way ANOVA. LTP was induced in both genotypes (*; +/+: p<0.0001, N = 16; –/–: p=0.0125, N = 16) with no significant difference between genotypes. C. Basal synaptic transmission in awake, behaving mice is reduced in Shank2^−/–^ mice compared to wildtype controls. *: p=0.011 (difference between genotypes in a two-way ANOVA; N_+/+_ = 12, N_–/–_ = 8).

How can we consolidate the discrepancy between reduced synaptic transmission and increased LTP? Is there a mechanistic explanation linking these two findings? In Shank3^-/-^ mice, reduced LTP has been associated with NMDA as well as AMPA receptor hypofunction (Wang *et al.*, 2011; Kouser *et al.*, 2013). Both mechanisms seem plausible, since hippocampal CA1 LTP is dependent on both NMDA and AMPA receptors in its induction and expression, respectively (Nicoll, 2017). Shank2^-/-^ mice show reduced AMPA receptor dependent basal synaptic transmission (Figure 1C), and, in certain conditions, a reduction in AMPA/NMDA receptor ratios (Figure 1D). We thus set out to test whether synapses lacking SHANK2 might possess fewer AMPA receptors or be functionally silent (i.e. lack them altogether), which would render these synapses salient substrates for the expression of LTP. Indeed, minimal stimulation revealed markedly higher failure rates in Shank2^-/-^ mice than wild-type controls at hyperpolarized potentials (-60 mV, when only AMPA receptor containing synapses contribute to EPSCs) compared to depolarized potentials (+40 mV, when synapses lacking AMPA but possessing NMDA receptors can also pass currents) (Figure 3A). Given that the apparent synaptic potency was not different between genotypes (Figure 3B), this suggests that the reduced AMPA/NMDA receptor ratios (Figure 3C) are a direct consequence of an increased fraction of silent synapses in Shank2^-/-^ mice (Figure 3D). At the same time, these silent synapses might provide the structural framework that facilitates the expression of LTP in Shank2^-/-^ mice.

**Figure 3.**
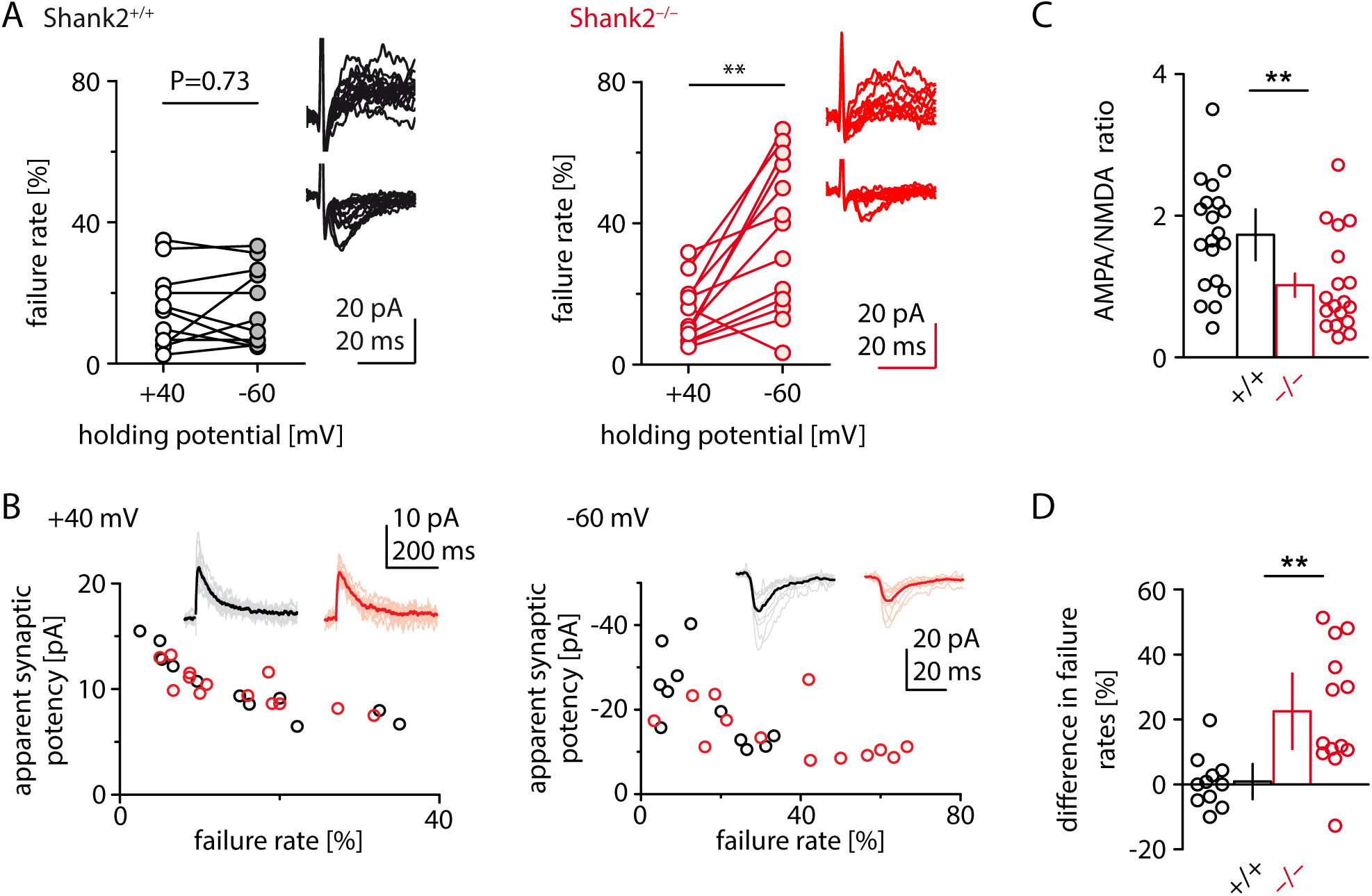
Minimal stimulation reveals insufficiently matured synapses in Shank2^-/-^ mice. A. EPSCs were recorded at different holding potentials under minimal stimulation *in vitro*. Failure rates indicate the proportion of AMPA receptor containing synapses (-60 mV) as a fraction of total synapses (+40 mV). Failure rates at different holding potentials are plotted for control (left) and Shank2^−/–^ mice (right).**: p=0.0012 (paired Student’s t-test; n(N)_+/+_ = 11(6), n(N)_–/–_ = 13(6)). Example traces at right (top panel: EPSPs recorded at +40 mV, bottom panel: EPSPs recorded at −60 mV). B. Synaptic potency as apparent in minimal stimulation experiments is not different between genotypes when differences in failure rates are taken into account, for both NMDA receptor mediated currents (+40 mV, left) and AMPA receptor mediated currents (-60 mV, right). Example traces at top. C. AMPA/NMDA receptor ratios calculated from average EPSP of minimal stimulation experiments are smaller in Shank2^-/-^ mice than in wildtype controls. **: p=0.005 (Student’s t-test; n(N)_+/+_ = 20(6), n(N)_–/–_ = 18(8)). D. The relative difference between failure rates at hyperpolarized versus depolarized potentials (F_-60_ minus F_+40_) is significantly higher in Shank2^-/-^ mice than in wildtype controls. **: p=0.0022 (Student’s t-test; n(N)_+/+_ = 11(6), n(N)_–/–_ = 13(6)).

A similar excess of silent synapses has been described in mice lacking Sapap3 (Wan, Feng and Calakos, 2011), a GKAP family protein that directly interacts with SHANKs (Boeckers *et al.*, 1999; Naisbitt *et al.*, 1999; Yao *et al.*, 1999) and whose absence in mice causes obsessive-compulsive behavioural traits (Welch *et al.*, 2007) – which are also typical in autistic individuals. Likewise, FMR1^-/-^ mice, which model the autism-related Fragile X syndrome, show altered plasticity and synapse maturation in the barrel cortex (Harlow *et al.*, 2010). In another mouse model for intellectual disability (SYNGAP1 haploinsufficiency) hippocampal synapses are unsilenced prematurely, adversely impacting learning and memory in the adult animal (Rumbaugh *et al.*, 2006; Clement *et al.*, 2012). These and the present study draw a picture of synapse maturation as a tightly controlled process, the dysregulation of which seems of relevance for a range of neurodevelopmental disorders; this process must therefore be investigated further in future studies, in particular with regard to how it can be influenced by therapeutic approaches for the potential treatment of autism and related disorders.

In summary, we consistently observe increased LTP in hippocampal Schaffer collateral-CA1 synapses of Shank2^-/-^ mice *in vitro*, as well as in awake behaving animals. We have uncovered a developmental synapse phenotype that could link the phenomena of decreased synaptic transmission and increased LTP in Shank2^-/-^ mice. These data add weight – and mechanistic insight – to our initial conclusion that SHANK2 loss of function results in *enhanced* synaptic plasticity, highlighting the necessity to caution against generalizing therapeutic approaches that rely on boosting LTP as a strategy to treat ASD phenotypes.

## Methods

*Shank2*^−/–^ mice were bred in a C57BL/6J background with a heterozygous breeding protocol. The study was carried out in accordance with the European Communities Council Directive of September 22, 2010 (2010/63/EU) for care of laboratory animals and after approval of the local ethics and/or animal welfare committees (Berlin animal experiment authorities and the animal welfare committee of the Charité Berlin (File reference: T100/03) and Landesamt für Naturschutz, Verbraucherschutz und Umweltschutz, Nordrhein Westfalen, respectively). Wild-type littermates were used as a control throughout and experimenters were blind to the genotype of the tested animals for data collection and analysis.

Hippocampal brain slices were prepared from animals of both sexes as described(Schmeisser* *et al.*, 2012). Briefly, mice were anesthetized with isoflurane and decapitated. Brains were rapidly removed and transferred to ice-cold ACSF slicing solution. Slicing ACSF contained (in mM): 87 NaCl, 26 NaHCO_3_, 50 sucrose, 25 glucose, 3 MgCl_2_, 2.5 KCl, 1.25 NaH_2_PO_4_, 0.5 CaCl_2_. Recording ACSF contained (in mM): 119 NaCl, 26 NaHCO_3_, 10 glucose, 2.5 KCl, 2.5 CaCl_2_, 1.3 MgCl_2_, 1 NaH_2_PO_4_. All ACSF was equilibrated with carbogen (95% O_2_, 5% CO_2_). Tissue blocks containing the hippocampus were mounted on a Vibratome (Leica VT1200) and cut into horizontal slices of 300 μm. For submerged slice storage (used for minimal stimulation experiments), slices were – after preparation – kept submerged in ACSF at 34°C for 30 min, then slowly cooled to room temperature where they were left to recover for at least 30 min up to 5 hours. Recordings were performed in slices submerged in ACSF and at room temperature. For a subset of experiments (Figure 1), we reproduced a range of storage and recording conditions from Won et al.(Won *et al.*, 2012): Immediately after preparation, slices were transferred to in an ACSF/oxygenated air interface chamber and allowed there to recover until recording, for at least one and at most five hours. Recordings were performed in a submerged recording chamber, storage and recording temperature for these experiments was 34°C. Mouse age for *in vitro* experiments was 8-9 weeks for experiments shown in Figure 1A and D, and 3-4 weeks for experiments in figure 3. For the remaining *in vitro* experiments (Figure 1B, C), mouse age is indicated in the figure legend.

For all *in vitro* experiments, data was recorded with an Axopatch 700A Amplifier, digitized at 5 kHz, filtered at 2 kHz and recorded in IGOR Pro 4.0. Evoked postsynaptic responses were induced by stimulating Schaffer collaterals in CA1 stratum radiatum. Field excitatory postsynaptic potentials (fEPSPs) were recorded in stratum radiatum. fEPSP rising slopes were fitted to 20 to 80% of the fEPSP amplitude. LTP was induced by a single tetanus of 100 pulses at 100 Hz. For whole-cell patch-clamp recordings, pipettes had resistances of 2–3 Mω. Liquid junction potential was not corrected. Series resistance (not compensated) was constantly monitored and was not allowed to increase beyond 22 Mω or change by more than 20 % during the experiment. Compound EPSCs were recorded at –60 and +40 mV with a cesium-based intracellular recording solution containing (in mM) 145 CsCl, 10 HEPES, 0.2 EGTA, 2 MgCl_2_, 2 NaATP, 0.5 NaGTP, 5 phosphocreatine (osmolarity 305 mOsm, pH adjusted to 7.2 with CsOH). The AMPA receptor mediated component of the EPSC was estimated by measuring the peak amplitude of the averaged EPSC at −60 mV. The NMDA receptor mediated component was estimated at +40 mV by measuring the amplitude of the averaged EPSC 25 ms after stimulation.

For minimal stimulation, stimulation frequency was 0.2 Hz and the stimulation electrode was placed to produce a single-peak response. At +40 mV holding potential, stimulation intensity was reduced until transmission failures were observed (in ∼ 10-40% of events), and 20-50 events were recorded. Cells were subsequently clamped to −60 mV holding potential, and 30-50 events were recorded at the same stimulation intensity. In a subset of experiments, this order was reversed, i.e. stimulation intensity was adjusted and miniature EPSCs (minEPSCs) were recorded at −60 mV first, before cells were clamped to +40 mV. We did not observe systematic differences in failure rates or amplitudes of minEPSCs between the two regimes. Experiments with linearly increasing or decreasing failure rates and/or minEPSC amplitudes at any holding potential were excluded from the analysis. Post-hoc analysis counted a failure at depolarized potentials whenever the EPSC charge 10 to 90 ms after stimulation did not exceed a fixed threshold of 0.15 pC. Failures at hyperpolarized potentials were defined as events with a minEPSC peak smaller than twice the signal noise of that recording (signal noise: standard deviation of the signal in a 3 ms time window averaged over all sweeps at a certain holding potential). Signal noise was not different between experimental groups, experiments with high background noise were excluded from the analysis. While the absolute failure rates depended on the criteria for failure vs successes were set, the relative difference between genotypes did not. Apparent synaptic potency was calculated as the EPSC amplitude of all events categorized as non-failures (at −60 mV by measuring the peak amplitude of the averaged EPSC, at +40 mV by measuring the amplitude 25 ms after stimulation). The mean stimulation intensity used to evoke near-unitary events was not significantly different between genotypes (Student’s t-test: P = 0.967; +/+: 0.62 ± 0.29 V, n(N) = 10(5); –/–: 0.65 ± 0.27 V, n(N) = 13(6)).

For *in vivo* experiments, male 7-8 week old mice were implanted (under anesthesia) with a stimulation electrode in the Schaffer collaterals (2.0 mm posterior and 2.0 mm lateral to Bregma) and a recording electrode in the Stratum radiatum of the dorsal CA1 region (1.9 mm posterior and 1.4 mm lateral to Bregma), as described previously (Kemp and Manahan-Vaughan, 2008). Animals recovered for ∼10 days before experiments were commenced. Before each experiment input/output (I/O) properties were recorded by increasing the stimulation intensity stepwise (Figure 2C). Experiments were carried out in recording chambers (20 (length) x 20 (width) x 30 (height) cm) where animals could move freely and had access to food and water *ad libitum.* During recordings, implanted electrodes were connected via a flexible cable and a rotatable commutator to the stimulation unit and amplifier. Test-pulse stimulation was set to elicit 40% of the maximal I/O response. Stimuli of 0.2 ms duration were applied at a frequency of 0.025 Hz and recorded with a sample rate of 10 kHz. To induce long-lasting LTP, two different induction protocols were used: Protocol 1 contained 4 trains of 50 pulses at 100 Hz with 5 min inter train interval (N_+/+_ = 8, N_–/–_ = 8). Protocol 2 contained 2 trains of 50 pulses at 200 Hz with 5 min inter train interval (N_+/+_ = 8, N_–/–_ = 7). Results from both protocols were quantitatively similar and pooled for Figure 2A. Short-lasting LTP was elicited using the following protocols: Protocol 3, a single train of 100 Hz stimulation (N_+/+_ = N_–/–_ = 8), and Protocol 4, two trains of 100 Hz stimulation with 5 min inter train interval (N_+/+_ = N_–/–_ = 8). Results from Protocol 3 and 4 were quantitatively similar and pooled for extended data Figure 2B. Statistical tests for expression of LTP within an experimental group involved comparing post-tetanus responses to pre-tetanus baseline with time as a continuous and pre/post induction as a categorical variable. Tests between genotypes compared post-induction time points only, with time as a continuous and genotype as a categorical variable.

Analyses were performed using custom written procedures in IGOR Pro and MATLAB. Data in graphs and in the text are presented as mean ± standard error unless indicated otherwise. Unpaired two-tailed Student‘s t-test (short: Student‘s t-test) and multiple way ANOVA were used to test for statistical significance. Results were considered significant at p < 0.05. Stimulus artifacts were blanked or cropped in sample traces. Sample sizes are given as number of experiments (n, applies to *in vitro* experiments only) and number of animals (N).

## Acknowledgements

This work was supported by the Cluster NeuroCure (Exc257), the Bernstein Center for Computational Neuroscience, the Einstein Foundation and the Deutsche Forschungsgemeinschaft (SFB665, SFB874/B1, SFB958, SFB1280/A4, SPP1665, GRK1123, SH650/2-1). We thank Susanne Rieckmann, Anke Schönherr, and Karin Bloch for assistance with mouse breeding, Dr. Christa Thöne-Reineke for overseeing animal transports, and Stephan Jansen for technical assistance with the *in vivo* experiments.

